# The growth benefits and toxicity of quinone synthesis are balanced by a dual regulatory mechanism and substrate limitations

**DOI:** 10.1101/2025.02.15.638467

**Authors:** Siliang Li, Jiangguo Zhang, Caroline M. Ajo-Franklin, Oleg A. Igoshin

**Author notes:** Co-corresponding authors (these authors supervised this work jointly and equally). These authors contributed equally (Siliang Li, Jiangguo Zhang).

## Abstract

Quinones play a central role in maintaining redox balance and conserving energy but can trigger oxidative stress at high levels. However, the mechanisms by which microbes regulate quinone levels remain poorly0020understood, hindering effective metabolic engineering to modulate microbes for quinone production. Here, we show that the synthesis of the menaquinone precursor DHNA in the lactic acid bacterium *Lactococcus lactis* is regulated by a combined genetic, enzymatic, and metabolic mechanism. Using synthetic biology approaches, we found that enzymes MenF and MenD both contribute to DHNA regulation, with MenD playing a more prominent role in controlling DHNA concentrations. A mathematical model elucidates a two-phase regulatory pattern resulting from the interplay of reversible flux and allosteric feedback inhibition, where either MenF or MenD can serve as the regulatory enzyme, depending on their relative expression ratio. Additionally, the overproduction of DHNA is constrained by substrate availability, ensuring a sufficient but not excessive DHNA level to benefit cell growth while mitigating cytotoxicity. Collectively, these mechanisms maintain a fine-tuned physiological quinone level and suggest that modulating substrate supplement and MenF-to-MenD ratio could be keys for engineering DHNA production

## Introduction

Quinones, such as menaquinone (MK, vitamin K2) and its precursor 1,4-dihydroxy-2-naphthoic acid (DHNA), are important microbial metabolites that support electron transport in respiratory growth(1–5), regulate redox stress(1, 6, 7), and enhance human cardiovascular and bone health(8). Diverse microbes produce rich amounts of quinones in fermented food, such as cheese, natto, and fermented milk(9, 10). There are particular interests in optimizing microbial production of MK and DHNA to enhance dietary quinone content and improve the nutritional value of fermented food(11, 12).

However, metabolic strategies to engineer MK and DHNA production have yielded contradictory results(11), largely due to limited understanding of the regulatory mechanisms governing quinone synthesis. MK biosynthesis starts from chorismate and proceeds through a seven-enzyme pathway consisting of MenF, MenD, MenH, MenC, MenE, MenB, and MenI to produce DHNA. Subsequently, MenA joins DHNA and prenyl diphosphate to produce demethylmenaquinone (DMK), and MenG demethylates DMK to generate MK (**Fig. 1A**). Strategies have attempted to enhance DHNA supply by overexpressing individual enzymes(13–17) or the entire DHNA synthesis operon(14, 18). However, these efforts often lead to inconsistent and modest increases in DHNA/MK levels despite upregulated transcription of the respective genes(14, 15). Conflicting outcomes have also been reported across different studies; for example, overexpressing MenF or MenD enhanced MK production in some cases but not others(13, 15–17). These data suggest that a regulatory network might exist to control DHNA and MK levels and buffer against genetic perturbations. While these regulations are physiologically plausible, as microbes must resist quinone overproduction to avoid oxidative stress and growth inhibition(4), the molecular mechanisms remain elusive.

**Figure 1:**
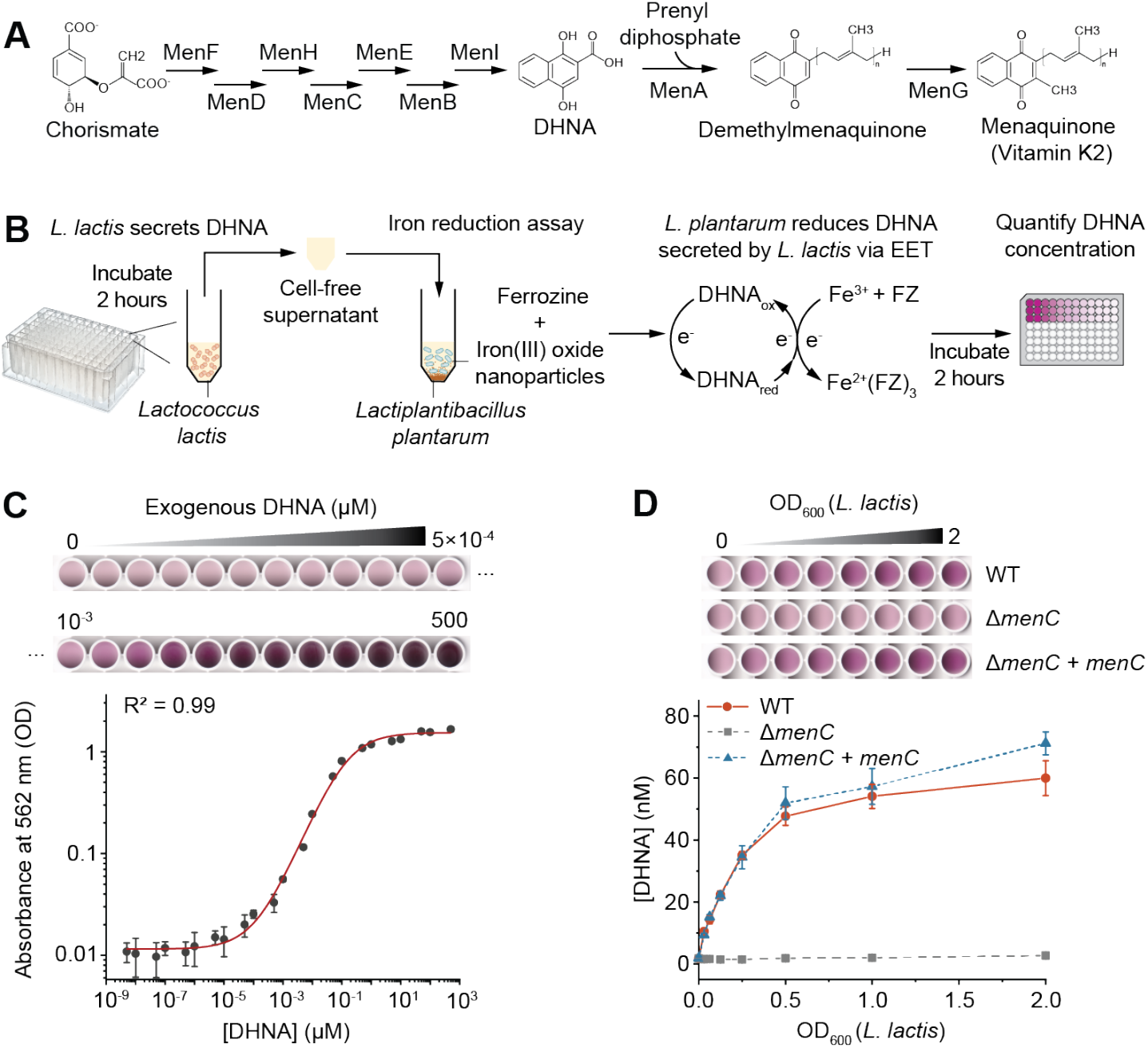
A biosensing system for quantitative detection of DHNA at picomolar concentration. **(A)** Schematic showing the menaquinone biosynthesis pathway. DHNA is the precursor of menaquinone. **(B)** Workflow of the DHNA biosensing system. *L. lactis* was incubated for 2 hours to allow DHNA synthesis. The cell-free supernatant was collected and mixed with *L. plantarum* and iron reduction reagents, including iron(III) oxide and ferrozine (FZ). *L. plantarum* utilizes DHNA secreted by *L. lactis* to reduce Fe^3+^ to Fe^2+^. The resulting Fe^2+^ reacts with ferrozine, producing a magenta-color product whose absorbance correlates with DHNA concentration. **(C)** Calibration curve of DHNA concentration versus absorbance at 562 nm. *L. plantarum* was incubated with varying concentrations of exogenous DHNA. The [DHNA]-OD_562_ curve was fitted to a non-linear function. The top panel shows a scanned image of the assay plate. **(D)** *L. plantarum* can detect DHNA secreted from varying cell densities of quinone-producing *L. lactis* but not from the deficient mutant. Δ*menC + menC* represents *menC* gene complementation under a constitutive promoter on a plasmid in the Δ*menC* background. DHNA concentrations were calculated based on the calibration curve in **(C)**. The top panel shows a scanned image of the assay plate. All data represent mean ± 1 s.d. for three biological replicates.

Previous studies shed light on the quinone regulatory mechanism, but a systems-level understanding has not been achieved. Studies have revealed that MenD can be allosterically inhibited by DHNA in *Staphylococcus aureus*(19) and *Mycobacterium tuberculosis*(20), suggesting an end-product feedback mechanism. In addition, MenF is also known to be capable of catalyzing both forward and reverse conversion of chorismate to isochorismate(21). However, it is unclear how these individual mechanisms interactively act to control DHNA production.

Here, we study one of the major quinone producers, *Lactococcus lactis*, and reveal a multi-layer mechanism regulating DHNA synthesis. Leveraging a novel biosensing system, we quantified DHNA concentration at picomolar sensitivity. We then synthetically perturbed enzyme levels on genetic constructs and found that DHNA concentration can be modulated by both MenF and MenD but to different degrees. A mathematical model uncovers a two-phase regulatory pattern resulting from the interplay of reversible flux and end-product feedback inhibition, where the regulatory roles of MenF and MenD shift based on their relative expression levels. The model also predicts limited availability of the substrate chorismate, which restrains DHNA from being overproduced. These mechanisms can collectively contribute to quinone homeostasis and imply new strategies to efficiently engineer quinone production.

## Results

### A biosensing system allows sensitive and quantitative DHNA detection

Since DHNA is the common precursor for all MK derivatives, we sought to interrogate the regulatory mechanism of quinone synthesis by focusing on DHNA. To precisely measure DHNA concentration, we constructed a *L. lactis* mutant lacking DHNA-utilizing pathways. We knocked out *menA* (encodes for DHNA prenyltransferase) to prevent DHNA from being utilized for DMK and MK synthesis (**Fig. 1A**). Additionally, we knocked out *noxAB* (encodes for NADH-quinone oxidoreductases) to prevent DHNA from interacting with quinone reductases(22), which may otherwise alter DHNA redox state and interfere with DHNA quantification.

We next developed a high-throughput method for quantitative DHNA detection. While previous studies mainly measured DHNA concentration with high-performance liquid chromatography (HPLC)(23– 26), this approach requires complex sample preparation and purification processes and has a limited detection of 1 µM DHNA(26). We previously discovered that the quinone auxotroph *Lactiplantibacillus plantarum* can utilize nanomolar DHNA for extracellular electron transfer (EET) and produce quantitative signals through the colorimetric iron reduction assay(27–29). Based on this phenomenon, we used *L. plantarum* as a biosensor to detect DHNA from *L. lactis* (**Fig. 1B**). This method mixes the cell-free supernatant (CFS) of *L. lactis* with sensor *L. plantarum*, iron(III) oxide, and the colorimetric reagent ferrozine (**Fig. 1B**). When CFS contains DHNA, *L. plantarum* uses DHNA to reduce Fe^3+^ to Fe^2+^ through extracellular electron transfer, and Fe^2+^ subsequently reacts with ferrozine to form a magenta-colored product that can be quantified at 562 nm (**Fig. 1B**).

We first established a calibration curve for DHNA quantification using the biosensing system. We exposed *L. plantarum* to varying concentrations of exogenous DHNA ranging from 5×10^−9^ to 5×10^2^ µM. A quantitative correlation was observed between the absorbance at 562 nm and the DHNA concentrations (**Fig. 1C**). Notably, the sensor was able to detect DHNA as low as 100 pM, demonstrating a sensitivity 10^4^ times greater than HPLC(26). We then fitted the experimental data using a non-linear function to obtain the OD_562_-[DHNA] calibration curve (**Materials and Methods**). This calibration curve indicates our ability to reliably detect DHNA concentrations within a linear range of 100 pM to 1 µM.

To evaluate the biosensor’s ability to detect DHNA secreted by *L. lactis*, we collected CFS from *L. lactis* strains with varying cell densities (OD_600_). These include strains with a wild-type (WT) DHNA pathway, a deficient DHNA pathway (Δ*menC*), or a complemented pathway with *menC* expressed under a constitutive promoter (Δ*menC + menC*). The biosensor detected no DHNA from the Δ*menC* strain (**Fig. 1D**). In comparison, DHNA was detected from strains containing the WT or *menC*-complemented DHNA pathway, with DHNA concentrations (10-60 nM) proportional to *L. lactis* cell density (OD_600_=0.03-2) (**Fig. 1D**). These DHNA concentrations fall in the middle of the linear range of the calibration curve, demonstrating reliable detection. These results confirmed that the biosensing system could specifically and reliably detect DHNA from CFS of *L. lactis*. We also noticed a trend of DHNA saturation at higher cell densities of *L. lactis* (OD_600_ =0.5-2.0). To avoid potential inhibition of DHNA synthesis at high cell densities, we decided to use *L. lactis* cultures with an OD_600_ of 0.2 for subsequent experiments to explore the regulatory mechanism of DHNA synthesis.

### DHNA synthesis exhibits different degrees of sensitivity to MenF or MenD

We next sought to use synthetic biology to alter the expression levels of the enzymes and explore their regulatory roles in DHNA synthesis. We focused on three enzymes in the DHNA synthesis pathway: MenC, MenF, and MenD. The o-succinylbenzoate synthase MenC catalyzes one of the intermediate reactions, whereas the isochorismate synthase MenF and the SEPHCHC synthase MenD catalyze the first and the second reaction, respectively (**Fig. 1A**). In bacteria such as *Escherichia coli*, MenF’s product isochorismate also participates in siderophore synthesis, and MenD is considered the first step solely committed to DHNA synthesis(30). However, *L. lactis* lacks the pathway for siderophore synthesis, and we consider MenF as the first step in *L. lactis* for DHNA synthesis.

To perturb the enzyme levels, we used combined transcriptional and translational strategies. For transcriptional regulation, we deleted the respective genes from the *L. lactis* genome and controlled their expression on a plasmid from a promoter that can be induced by the antimicrobial peptide nisin(31) (**Fig. 2A**). For translational regulation, we used either strong or weak ribosome binding sites (RBSs) to control the translation initiation rate (**Fig. 2A**). Additionally, we fused mCherry to the N-terminal of MenC, MenF, and MenD, and measured fluorescent intensity to monitor enzyme expression levels. Notably, *menF* and *menD* knockouts were constructed in the Δ*menC* background. In these cases, *menC* was expressed on the plasmid under a constitutive promoter, which yielded a DHNA level similar to the wild-type pathway (**Fig. 1D**), ensuring the effects of MenF or MenD on DHNA could be studied without interference.

**Figure 2:**
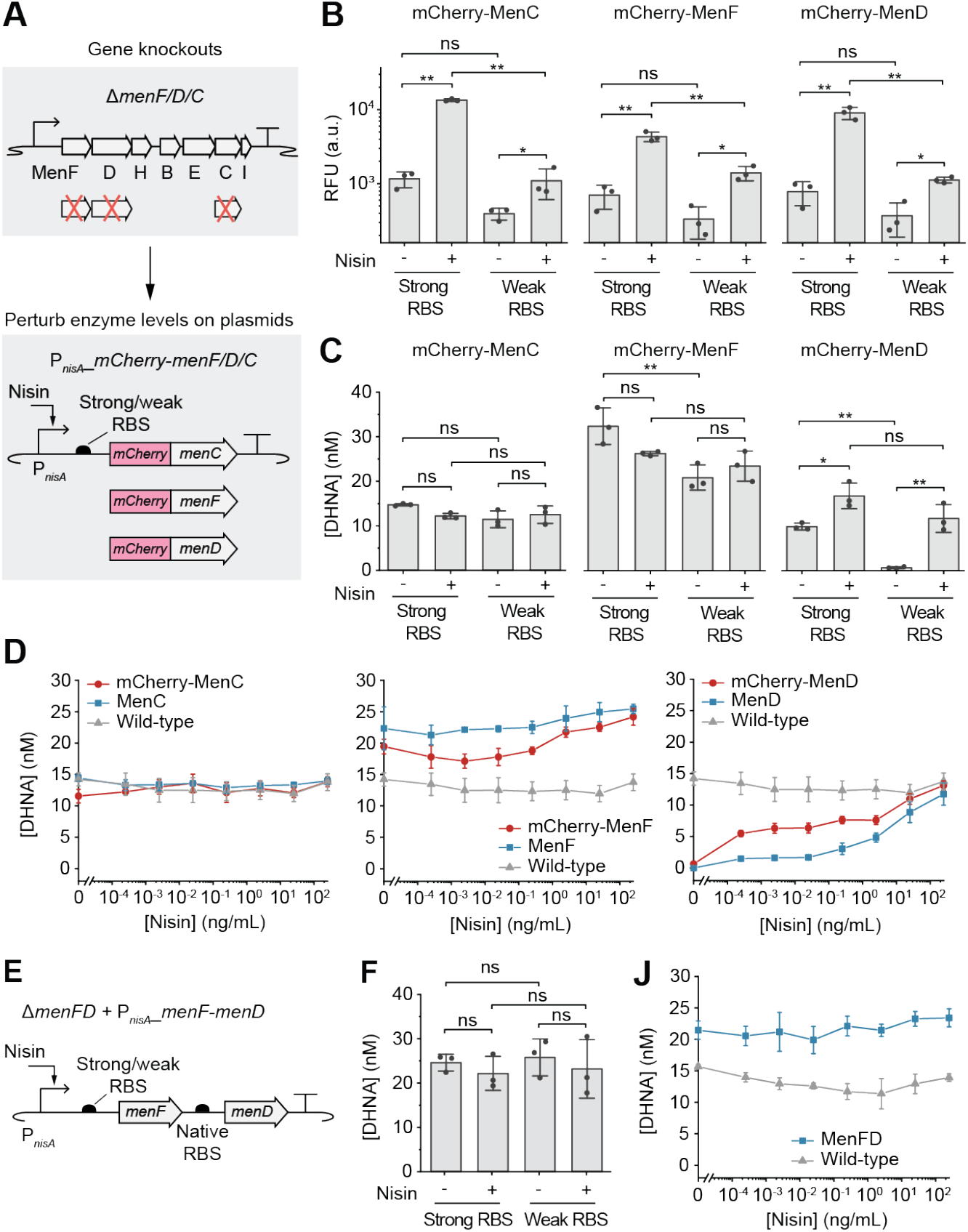
Effects of enzyme level perturbation of MenC, MenF, MenD, and MenFD on DHNA synthesis. **(A)** The *menC, menF*, or *menD* genes were knocked out from the genome and reintroduced under the control of a nisin-inducible promoter (P_*nisA*_) on a plasmid with either a strong or a weak ribosome binding site (RBS). mCherry was fused to the N-terminal of each enzyme to monitor enzyme expression. **(B)** Enzyme expression levels with or without nisin induction (25 ng/mL) were quantified by the fluorescent intensity. RFU, relative fluorescent unit. **(C)** DHNA concentrations under varying enzyme expression levels. **(D)** DHNA concentrations from perturbations of mCherry-tagged MenC, MenF, or MenD were compared to those resulting from the untagged enzymes and the wild-type pathway. All genes were expressed under the weak RBS. Varying concentrations of nisin were used to induce different enzyme expression levels. Nisin was also added to the wild-type pathway as a control. **(E)** The *menFD* genes were knocked out from the genome, and the identical gene cassette was expressed under P_*nisA*_ with a strong or weak RBS. **(F)** DHNA concentrations under varying MenFD expression levels, with or without nisin induction (25 ng/mL). **(J)** DHNA concentrations resulting from MenFD perturbations were compared to the wild-type pathway. All data represent mean ± 1 s.d. of n=3 biological replicates. P-values were determined by one-way ANOVA with Tukey’s test. *p < 0.05, **p < 0.01, ***p < 0.001, ns, not significant.

When nisin was added to induce enzyme expression with the strong RBS, we observed 11.6, 6.2, and 11.6-fold increases in fluorescence for mCherry-tagged MenC, MenF, and MenD, respectively (**Fig. 2B**). With the weak RBS, fluorescence showed 2.8, 4.2, and 3-fold increases for mCherry-tagged MenC, MenF, and MenD, respectively (**Fig. 2B**). These results indicate that our constructs can effectively modulate enzyme levels. However, these enzyme level perturbations did not correlate with changes in DHNA concentrations. The basal expression level of mCherry-tagged MenC or MenF (without induce) was already sufficient to drive DHNA synthesis, and nisin-induced overexpression did not further increase DHNA concentrations (**Fig. 2C**). Only upon induction of mCherry-MenD did the DHNA concentration increase by 1.7 and 17.5-fold with the strong and weak RBS, respectively (**Fig. 2C**). These results indicate that DHNA synthesis is sensitive to the perturbation of mCherry-MenD but not to mCherry-MenC/MenF.

To validate these findings, we removed the mCherry tags and perturbed the untagged enzymes using the nisin-inducible promoter and the weak RBS. The results mirrored those of the tagged enzymes: DHNA concentration was regulated only by different MenD levels but not by MenC or MenF (**Fig. 2D**). Additionally, we compared the DHNA concentration to the wild-type pathway and found that perturbations of MenC and MenD did not yield DHNA levels exceeding the wild-type (**Fig. 2D**). Interestingly, basal MenF expression on the plasmid produced a DHNA level 1.5-fold higher than the wild-type, but further increase in MenF expression could not overproduce DHNA (**Fig. 2D**). These data suggest that DHNA synthesis can also be modulated by MenF; however, DHNA concentration is plateaued at the basal expression level, resisting overproduction.

We speculated that the inability to further increase DHNA production by overexpressing MenF might result from a flux limitation imposed by genomic MenD expression level. To test this hypothesis, we knocked out the adjacent *menF* and *menD* genes from the genome and co-regulated their expression on the plasmid (**Fig. 2E**). Interestingly, while MenD alone could regulate DHNA levels (**Fig. 2D**), simultaneous perturbation of MenFD did not result in significant changes in DHNA concentrations (**Fig. 2F**). Although DHNA level was elevated compared to the wild-type cells, further increase in MenFD expression did not overproduce DHNA (**Fig. 2J**). Thus, these results indicate that the saturation in DHNA production is not due to the limited MenD expression. Surprisingly, these data also show that the combined perturbation of MenFD suppresses MenD’s regulatory influence on DHNA.

### Interplay of reversible flux, allosteric feedback inhibition, and substrate limitation regulates DHNA synthesis

Based on the experimental data, we aimed to explain why DHNA exhibits varying sensitivities to MenF and MenD and why its overproduction is resisted. To this end, we have constructed a mathematical model of the pathway that includes two critical features extracted from the literature: DHNA allosterically inhibits MenD(19, 20, 32), and MenF catalyzes both the forward and reverse reactions(21). We then used this model to perform a steady-state kinetic analysis and investigate how changes in enzyme levels would affect DHNA concentrations (**Materials and Methods**).

To develop this model (**Fig. 3A**), we initially assumed the concentration of chorismite, the first precursor and MenF’s substrate, is constant. We hypothesized this is a reasonable assumption because chorismate is the key branch point for many aromatic compounds(33), and cells would supply and use chorismate at a constant rate that is unlikely to be affected by our genetic perturbations. The model then considers the reversible conversion of chorismate to isochorismate catalyzed by MenF (**Equation 2**), as well as DHNA’s non-competitive inhibition on MenD (**Equation 3**). The remaining reactions follow irreversible Michaelis-Menten kinetics (**Equations 7-11**). Given that *menA* is knocked out and the downstream MK pathway no longer utilizes DHNA, we modeled the DHNA outflux due to its passive diffusion to the extracellular space (**Fig. 3A**). As experiments confirmed a constant DHNA accumulation rate in the supernatant over time (**Fig. S1**), we modeled DHNA outflux as a first-order process with the rate *k*_*DHNA*_(**Equations 11-12**).

**Figure 3:**
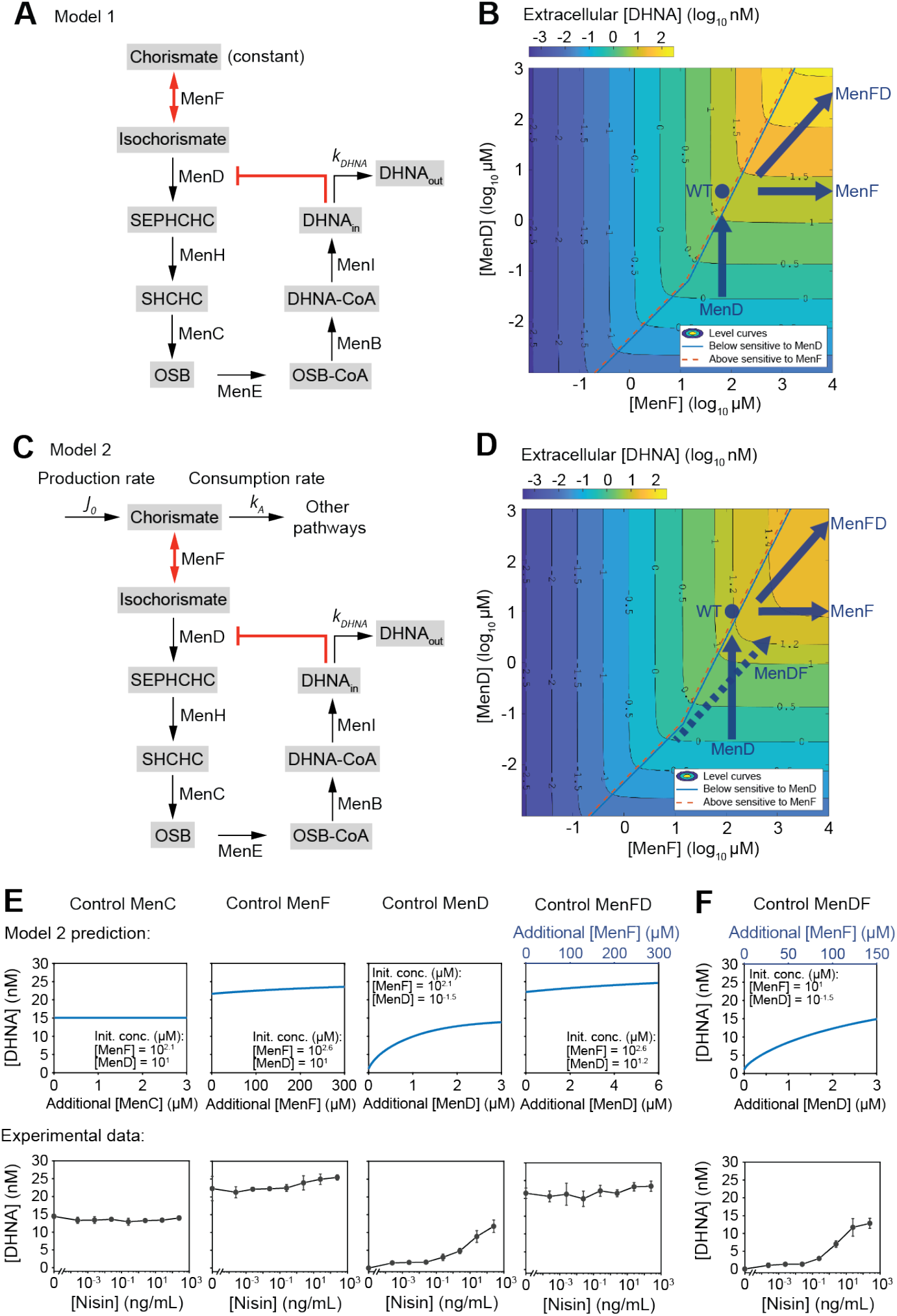
Steady-state kinetic models reveal that DHNA synthesis is regulated by a dual regulatory mechanism and constrained by substrate availability. **(A)** Model 1 considers a constant chorismite supply, DHNA outflux, and two regulatory mechanisms: MenF catalyzes a reversible reaction, and MenD is allosterically inhibited by DHNA. DHNA_in_, intracellular DHNA; DHNA_out_, DHNA diffused out of the cells. **(B)** Two-dimensional level curves derived from the numerical simulation of model 1. The level curves illustrate a two-phase regulatory pattern: DHNA can be regulated by either MenF (above the phase boundary) or MenD (below the phase boundary), depending on the relative MenF-to-MenD concentration ratios. The blue arrows predict changes in DHNA levels in response to the perturbations of MenF, MenD, and MenFD. The blue dot predicts the DHNA level of the wild-type pathway. **(C)** The revised model 2 considers chorismite production and consumption, DHNA outflux, and the two regulatory mechanisms. **(D)** The level curves derived from model 2 illustrate a third saturation phase, where DHNA synthesis is limited by the substrate chorismate and is no longer sensitive to enzyme perturbations. The dashed arrow predicts that DHNA is also sensitive to MenDF (the spatial swap of *menF* and *menD* gene positions). **(E)** Comparison between model-predicted and experimentally measured DHNA concentration in response to the perturbations of MenC, MenF, MenD, and MenFD. **(F)** Model prediction aligns with experimental data and confirms that DHNA can be regulated by MenDF. All genes were expressed under the nisin inducible promoter (P_*nisA*_) + weak RBS with the corresponding genes knocked out from the genome. For controlling MenFD and MenDF, the native RBS was used between *menF(D)* and *menD(F)*. All experimental data represent mean ± 1 s.d. of n=3 biological replicates.

Our model predicted that the steady-state DHNA concentration does not depend on the levels of MenH, MenC, MenE, MenB, or MenI, as perturbations of these enzymes can be balanced out by corresponding changes in their substrate concentrations (**Equations 7-11**). Importantly, the model suggests that DHNA can be regulated by both MenF and MenD (**Equation 13**), with the predominant regulatory enzyme determined by the MenF-to-MenD ratio and the DHNA concentration (**Equations 14-23**). To simulate this regulatory interplay, we performed a numerical simulation and generated a two-dimensional phase plot, i.e. level curves to predict the extracellular DHNA concentration under varying levels of MenF and MenD (**Fig. 3B**). The results revealed two phases in which either MenF or MenD acts as the rate-limiting enzyme. When MenF level is higher than MenD by approximately 100-fold (when [DHNA] < inhibition constant) or 10-fold (when [DHNA] > inhibition constant), MenF produces sufficient isochorismate, but the reaction flow is constrained by MenD, making MenD the rate-limiting step (**Fig. 3B**, below the phase boundary). On the flip side, when MenF level is lower than MenD by the same folds, the insufficient isochorismate production makes MenF the rate-limiting step (**Fig. 3B**, above the phase boundary). Critically, the allosteric inhibition of MenD by DHNA reduces MenD’s regulatory efficiency, making DHNA approximately 10 times less sensitive to the perturbation of MenD (**Fig. 3B**, turning point at the middle of the phase boundary).

Using the two-dimensional phase plot, we qualitatively estimated the potential regulatory range of MenF and MenD to interpret the experimental data. We speculated that the expression ratio of MenF and MenD should lie below the phase boundary, where DHNA is predominately regulated by MenD (**Fig. 3B**, blue arrows). We estimated the initial MenF or MenD levels based on the experimentally measured DHNA concentrations (**Fig. 2D, J**), and plotted the predictive dose-response curves of DHNA concentrations as a function of additional enzyme levels (**Fig. S1**). The model aligned with experimental observations that DHNA is sensitive to MenD perturbations but not to MenC and MenF (**Fig. S1**). However, the model also predicted that co-regulated increases in MenFD should elevate DHNA concentrations (**Fig. S1**), as increased MenD levels compensate for the increased MenF to allow greater reaction flux (**Fig. 3B**, top right blue arrow). This is inconsistent with experimental results, where DHNA concentrations remained unchanged despite increased MenFD levels (**Fig. 2J**).

We hypothesized that this inconsistent prediction for MenFD might be due to the model’s inability to account for saturation in DHNA synthesis: with an increase of both MenF and MenD, the flux continues to increase, and nothing can limit it. This is because the model assumed the substrate chorismate is constant and never limited. In contrast, the limited supply of the substrate chorismate could restrain the flux. To test this, we expanded the model to allow chorismate to be flux-limiting by including a production rate *J*_0_ and a consumption rate (by other cellular processes) of *k*_*A*_ (**Fig. 3C** and **Equations 24-25**). The revised model suggests that a limited supply of chorismate could preset a maximum DHNA concentration (**Equation 26**). In other words, chorismate production rate *J*_0_ sets the absolute limit to DHNA production flux. When the basal levels of the co-regulated MenF and MenD are high enough to make chorismate the rate-limiting factor, further increases in MenFD would no longer elevate DHNA (**Fig. 3D**, top right blue arrow). With these assumed enzyme level regimes, we obtained model predictions agreeing with experimental data (**Fig. 3E**). Thus, the model explains the effects of enzyme level perturbations on DHNA synthesis and predicts that the supply of chorismate is limited.

In addition to successfully explaining experimental data, the model makes two other intriguing predictions. First, to explain why MenF and MenFD overexpression could only increase the wide-type DHNA level by about 1.5-fold (**Fig. 2D, J**), the model predicts that the wild-type DHNA level is already near saturation (**Fig 3D**, blue dot). This high wild-type DHNA concentration, along with the limited supply of chorismate, can be used as a mechanism to restrain DHNA overproduction. Second, to explain why MenD resulted in a lower initial DHNA concentration than MenFD in the absence of nisin induction (**Fig. 2D, J**), the model implies that the basal expression level of MenD is lower when it is regulated alone compared to as part of the MenFD context (**Fig. 3D**). This means that the basal expression of the *menD* gene is reduced when positioned as the first gene downstream of the promoter, compared to when it is positioned after the *menF* gene. This is not surprising because gene arrangement in genetic constructs and operons can alter individual gene expression levels(34, 35). We hypothesized that if *menF* is positioned after *menD*, MenF would have a reduced basal expression level, which would allow the co-regulated “MenDF” to perturb DHNA concentrations (**Fig 3D**, dashed arrow, and **Fig. 3F**). Indeed, when we tested the plasmid with flipped *menF* and *menD* positions, the increased MenDF expression levels elevated DHNA concentrations (**Fig. 3F**). This suggests that the spatial gene position of *menF* and *menD* also plays a crucial role in regulating DHNA synthesis.

### Robust DHNA synthesis mitigates DHNA-induced toxicity and benefits *L. lactis* growth

As the model suggests that the DHNA in *L. lactis* is produced at a concentration close to saturation and overproduction is restrained by substrate limitations, we sought to understand the physiological implication of such a regulatory mechanism. Although *L. lactis* primarily relies on fermentation for energy conservation, studies have shown that *L. lactis* can also perform aerobic or anaerobic respiration using quinones, such as DHNA, DMK, or MK, as electron mediators(36–38). These respiratory processes, while not essential for*L. lactis* growth, can facilitate NAD^+^ regeneration by shuttling electrons from the NADH pool to external terminal electron acceptors, such as iron(36, 37) or copper(37). However, high quinone concentrations can induce oxidative stress and hinder cell growth(4). Thus, we hypothesized that such a regulatory mechanism is evolved to maintain DHNA homeostasis and balance the beneficial and adverse effects.

To examine the impact of excessive DHNA on *L. lactis* growth, we exposed wild-type *L. lactis* to varying concentrations (24 nM-100 µM) of exogenous DHNA and monitored the cell growth curves. We observed that additional DHNA greater than 98 nM caused a delay in exponential growth, and 100 µM DHNA reduced the final biomass by half (**Fig 4A, left panel**). The degree of decrease in specific growth rate was proportional to the excessive DHNA concentration (**Fig 4A, right panel**). These results indicate that higher-than-physiological DHNA concentration can impede *L. lactis* growth.

**Figure 4:**
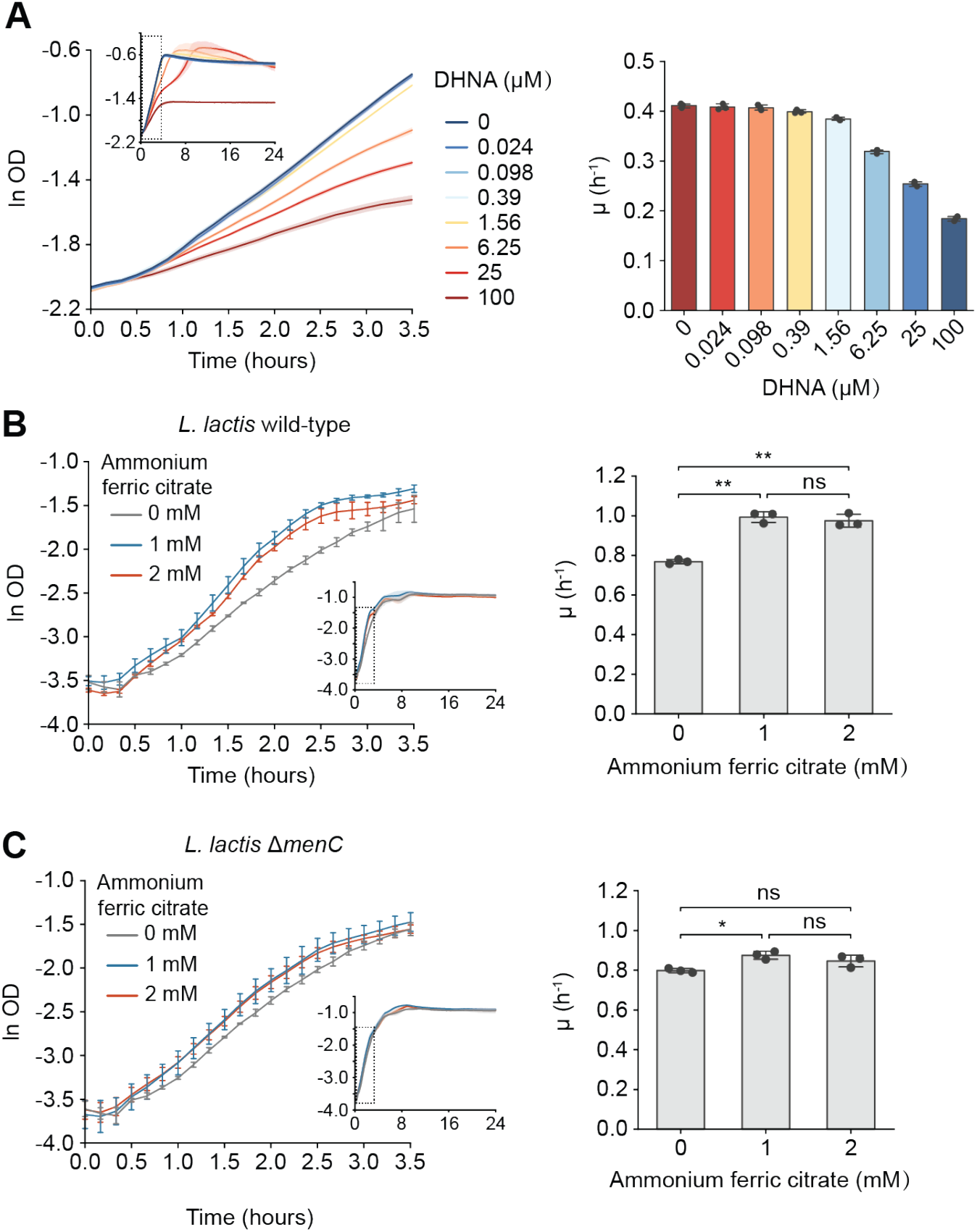
Robust DHNA synthesis mitigates DHNA-induced toxicity and promotes cell growth in the presence of terminal electron acceptors. **(A)** Growth curves and specific growth rate (μ) of wild-type *L. lactis* in response to different concentrations of exogenous DHNA. **(B)** Growth curves and specific growth rate (μ) of wild-type *L. lactis* in the presence of 1 or 2 mM ammonium ferric citrate under anaerobic conditions. **(C)** Growth curves and specific growth rate (μ) of *L. lactis* Δ*menC* in the presence of 1 or 2 mM ammonium ferric citrate under anaerobic conditions. The specific growth rate was determined by conducting the linear regression on the growth curve between 0.5 and 2.5 hours. All data represent mean ± 1 s.d. of n=3 biological replicates. P-values were determined by one-way ANOVA with Tukey’s test. *p ≤ 0.05, **p ≤ 0.01, ns, not significant.

We next investigated whether DHNA provides survival benefits to *L. lactis*. A previous study used an NAD^+^ regeneration-blocked *L. lactis* mutant strain and demonstrated that DHNA, but not MK, can mediate electron transfer from NoxAB to the terminal electron acceptor ferricyanide, restoring *L. lactis* growth under anaerobic conditions(36). To determine if DHNA could similarly benefit wild-type *L. lactis*, we inoculated wild-type *L. lactis* or the Δ*menC* mutant (*menA* and *noxAB* remained intact in the genome) with a terminal electron acceptor and monitored their growth under anaerobic conditions. To avoid the high toxicity of ferricyanide, we used ammonium ferric citrate (AFC), a less toxic alternative, as the terminal electron acceptor. When supplied with 1 or 2 mM AFC, *L. lactis* exhibited a 1.29 or 1.27-fold increase in growth rate during the exponential growth phase (**Fig 4B)**. This effect was significantly reduced in the DHNA synthesis deficient Δ*menC* mutant (**Fig 4C)**. These results confirm that DHNA synthesis offers additional growth advantages in the presence of terminal electron acceptors. We noted no difference in the growth rate for 1 or 2 mM AFC; this might be because AFC was overabundant. Additionally, a slight increase in growth rate was observed even in the Δ*menC* mutant (**Fig 4C)**, which we attributed to the exogenous redox-active molecules contained in the culture media, such as riboflavin from yeast extract, that may also serve as electron mediators to benefit *L. lactis* growth. These results support our hypothesis that maintaining a relatively high but not excessive DHNA level in *L. lactis* is critical to balance its benefits and adverse effects, highlighting the evolutionary advantage of the regulatory mechanism to control DHNA synthesis.

## Discussion

In this study, we revealed that DHNA synthesis in *L. lactis* is regulated through the interplay of the reversible flux, allosteric feedback inhibition, substrate limitation, and spatial gene arrangement. We found that DHNA synthesis can be primarily modulated by MenD, partially by MenF, but not by MenC. Additionally, a combined regulation of MenF and MenD could modulate DHNA synthesis only if the *menD* gene is positioned before the *menF* gene, showing a critical role of spatial gene arrangement in regulating DHNA synthesis. The steady-state kinetic model suggests the rate-limiting enzyme can be shifted between MenF and MenD depending on their relative concentration. By rationally assuming the concentration ranges of MenF and MenD, the model accurately predicts DHNA levels observed experimentally. The model also predicts that the wild-type DHNA level is close to saturation and is limited by the substrate chorismate. This sufficient but not overproduced DHNA level provides growth benefits while avoiding toxicity.

Our study sheds light on the mechanism of maintaining quinone homeostasis. Similar to other redox-active molecules, such as flavin(39) and phenazine(40), quinone synthesis is highly regulated to maintain beneficial physiological levels while preventing the negative consequences of redox imbalance. In *L. lactis*, the near-saturated DHNA production offers additional respiratory growth by facilitating electron transport to external electron acceptors (**Fig. 4B, C**), which can be a mechanism to enhance *L. lactis*’s adaptation to diverse environments(41). The saturated DHNA concentration can be preset by the limited availability of chorismate and is likely enforced by the high expression levels of MenF and MenD in the native pathway (**Fig. 3D**). While the mechanism driving high enzyme expression remains to be explored, it can be associated with gene arrangement of *menF* and *menD* altering the operon transcription and translation(34), or transcriptional activation of the *men* promoter(37, 42, 43) by regulators such as cAMP receptor protein(43) or catabolite control protein CcpA(37). Moreover, the allosteric inhibition of MenD by DHNA also contributes to quinone homeostasis by reducing the impacts of enzyme level fluctuations on DHNA synthesis. Given that the DHNA synthesis pathway is evolutionarily conserved(32), these regulatory mechanisms may also be employed by other bacterial species to maintain quinone homeostasis.

Our findings also suggest caveats and strategies for metabolic engineering to modulate DHNA and MK production. First, the flux of DHNA synthesis could not be modulated by intermediate enzymes. This is echoed in previous unsuccessful attempts to increase MK production by overexpressing intermediate enzymes for DHNA synthesis in *Escherichia coli*(13) and *Bacillus subtilis*(14, 15). Second, the simultaneous control of multiple enzymes within the DHNA operon may not always be effective in boosting its synthesis, as the outcome can depend on gene arrangement and the availability of substrate. This may explain why simply replacing the native promoter of the *men* operon with a constitutive one did not increase MK levels in *B. subtilis*(14). Third, increasing chorismate supply can be a key strategy to enhance DHNA and MK production. Supporting this, overexpression of enzymes in the chorismate synthesis pathway has been shown to enhance MK production(15). Finally, MenD can be a crucial target to regulate DHNA and MK production, which has been demonstrated in *E. coli*(13) and *B. subtilis*(17). The regulation efficiency of MenD can be further improved by engineering DHNA-insensitive variants to eliminate allosteric inhibition(19, 20). Indeed, mutant strains of *B. subtlis*(44) and *Flavobacterium meningosepticum*(21) resistant to 1-hydroxy-2-naphthoate (HNA, a DHNA analog) showed higher MK production than their parental strains. Collectively, the mathematical model presented in this study provides a coherent framework that explains prior observations for manipulating DHNA/MK synthesis across many organisms. This systematic understanding of the DHNA regulatory mechanism will facilitate more efficient metabolic engineering of the quinone synthesis pathway.

## Materials and Methods

### Strains, plasmids, and culture conditions

A list of strain used in this study is provided in **Table S1**. *E. coli* 5-alpha or 10-beta were used for cloning (New England Biolabs). All the *L. lactis* mutants were derived from the *L. lactis* subsp. *lactis* KF147. *L. plantarum* NCIMB8826 Δ*dmkA* Δ*ndh1* was used to sense the DHNA secreted from the *L. lactis* mutants.

Gene complementation in *L. lactis* was performed based on a modified backbone derived from pECGMC3 (Addgene #75441). Plasmids were assembled by Golden Gate Assembly(46) and were introduced in *L. lactis* by electroporation. *L. lactis* strains carrying plasmids were grown in the presence of 10 µg/mL erythromycin.

All *E. coli* strains used for cloning were grown in Terrific Broth (Sigma) containing 10 g/L glycerol. Preceding each experiment, *L. lactis* strains were grown in M17 broth (HiMedia) containing 0.5% glucose (gM17) at 30 °C without shaking, and *L. plantarum* was grown in commercial MRS (HiMedia) at 37 °C without shaking. A day before the iron reduction assay, *L. lactis* strains were subcultured in M17 broth containing 1% (w/v) mannitol (mM17) at 30 °C without shaking, and *L. plantarum* was subcultured in a modified MRS containing 1% (w/v) mannitol (mMRS) (**Table S2**) at 37 °C without shaking. The chemically defined medium containing 1% (w/v) mannitol (mCDM) (**Table S3**) was used in the iron reduction assay. The iron reduction assay was conducted in the anaerobic chamber with a temperature maintained at 30 °C.

### *L. lactis* mutant construction

*L. lactis* KF147 mutants were constructed by using double-crossover homologous recombination with the suicide plasmid pRV300(45). All the gene deletions were in-frame and were partial deletions with a small sequence segment retained at both the 5’ and 3’ of the gene. This avoids the detrimental effects of gene deletion on the expression of other genes within an operon. The up and down homologous arms (about 600 base pairs each arm) were amplified from the genome of *L. lactis* KF147 and inserted into the NotI-EcoRI digested pRV300 by using Golden Gate assembly(46). The resulting plasmids were then introduced to *L. lactis* KF147 by electroporation. Erythromycin-resistant colonies were selected and verified for genomic integration of the suicide plasmid (the first crossover) by using colony PCR. One positive colony was subsequently inoculated in gM17 (0.5% glucose) without antibiotics and passaged daily at a ratio of 1:1000. Starting on day 6 of passage, cells were diluted 2 × 10^6^ times and spread on five gM17 agar plates without antibiotics. The other day, cells were replica plated on gM17 agar plates containing 5 µg/mL erythromycin. This process was repeated daily until erythromycin-sensitive colonies appeared (the second crossover). Gene deletion was then verified by colony PCR and DNA sequencing.

### Supernatant iron reduction assay for DHNA quantification

*L. lactis* mutants and *L. plantarum* Δ*dmkA* Δ*ndh1* were inoculated from the glycerol stocks in gM17 or MRS and grown overnight for 15 h. The next day, *L. plantarum* Δ*dmkA* Δ*ndh1* was subcultured (1:100 v/v) into mMRS at a desired volume. *L. lactis* was subcultured in mM17 with an initial OD_600_ = 0.1 in a 96-deep well plate (450 µL culture per well). For *L. lactis* mutants carrying the nisin-inducible plasmids, the indicated concentrations of nisin were added when cells reached the exponential growth phase (OD_600_ = 0.4-0.6). After 15 h of overnight growth, *L. plantarum* and *L. lactis* cells were pelleted at 4000 g, 4 °C, for 10 min and washed twice with 1× PBS. *L. plantarum* cells were resuspended in mCDM to OD_600_ = 2. *L. lactis* cells were resuspended and diluted in mCDM to OD_600_ = 0.2 and incubated at 30 °C for 2 h to allow DHNA secretion. After 2 h, *L. lactis* cells were pelleted, and an aliquot of 160 µL cell-free supernatant (CFS) was transferred into a new 96-deep well plate. The CFS was mixed with 32 µL of 20 mM iron(III) oxide nanoparticles (< 50 nm particle size, Sigma), 32 µL of 20 mM ferrozine (Thermo Scientific), and 224 µL of *L. plantarum* resuspension. The plate was covered with aluminum foil and incubated in an anaerobic chamber (Whitley A45 Workstation) at 30 °C with 150 rpm shaking. After 2 h, an aliquot of 100 µL supernatant was collected to measure the absorbance at 562 nm using a plate reader (Tecan Spark). The DHNA concentration in each well was calculated using the DHNA – OD_562_ calibration curve.

### DHNA - OD_562_ calibration curve

The DHNA – OD_562_ calibration curve was generated by adding varying concentrations of exogenous DHNA to *L. plantarum* Δ*dmkA* Δ*ndh1*. After overnight growth in mMRS, the *L. plantarum* cells were pelleted at 4000 g, 4 °C, for 10 min, washed twice with 1× PBS, and resuspended in mCDM to OD_600_ = 2. In a 96-deep well plate, DHNA was 10-fold serially diluted in mCDM from 500 to 10^−9^ µM. Each dilution (160 µL) was combined with 32 µL of 20 mM iron(III) oxide nanoparticles (< 50 nm particle size), 32 µL of 20 mM ferrozine, and 224 µL of *L. plantarum* resuspension (or 224 µL mCDM for media control). The plate was covered with aluminum foil and incubated in the anaerobic chamber (Whitley A45 Workstation) at 30 °C with 150 rpm shaking. After 2 h, an aliquot of 100 µL supernatant was collected to measure the absorbance at 562 nm using a plate reader (Tecan Spark). The OD_562_ of media control was subtracted from the OD_562_ of the samples containing *L. plantarum* cells. The calibration curve was established by fitting the DHNA - OD_562_ curve to the Hill function provided below:

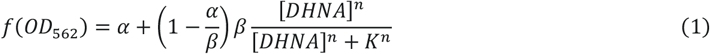

where α = 0.0115, β = 1.5338, K = 0.1330, n = 0.7102.

### *L. lactis* growth curve experiments

To examine the impact of exogenous DHNA on growth, the wild-type *L. lactis* KF147 was grown in gM17 overnight at 30 °C without shaking. The next day, cells were pelleted, washed twice with 1× PBS and resuspended with gM17 to OD_600_ = 1. A 10 mM DHNA stock solution was prepared in DMSO and was serially diluted in gM17. In a 96-well plate, 10 µL cell resuspension was combined with 90 µL gM17 containing different concentrations of DHNA. The final DHNA concentrations were 100, 25, 6.25, 1.56, 0.39, 0.098, 0.024 or 0 µM, with an initial cell OD_600_=0.1. The plate was placed in a humidity cassette and the OD_600_ was monitored using a plate reader (Tecan Spark) with the temperature maintained at 30 °C.

To examine the role of DHNA in anaerobic respiration, the wild-type *L. lactis* KF147 and the Δ*menC* mutant were grown in gM17 and subcultured in mM17 overnight at 30 °C without shaking. The next day, cells were pelleted, washed twice with 1× PBS and resuspended with gM17 to OD_600_ = 1. In the anaerobic chamber (Whitley A45 Workstation), 10 µL cell resuspension was combined with 85 µL mM17 + 5 µL 40 mM ammonium ferric citrate (AFC), 87.5 µL mM17 + 2.5 µL AFC, or 90 µL mM17. The final concentrations of AFC were 2, 1 or 0 mM, with an initial cell OD_600_=0.1. The plate was sealed with a transparent PCR plate seal (Axygen), and the OD_600_ was monitored using a Byonoy Absorbance 96 plate reader in the anaerobic chamber.

### Steady-state chemical kinetics analysis

#### 1. Model 1

To elucidate the rate-limiting step in the biosynthesis of 1,4-dihydroxy-2-naphthoic acid (DHNA) from chorismate in *L. lactis*, we employed a steady-state modeling approach using Michaelis-Menten kinetics. We simplified notation by assigning capital letters to intermediates in the pathway as follows: A for chorismate, B for isochorismate, C for SEPHCHC, D for SHCHC, E for OSB, F for OSB-CoA, G for DHNA-CoA. The enzyme MenF catalyzes a reversible reaction converting chorismate (A) to isochorismate (B)(43), and the isochorismate (B) production rate can be expressed as:

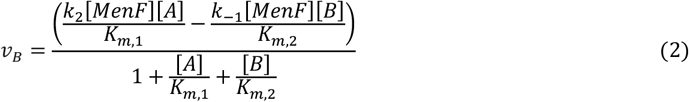

Subsequently, MenD facilitates the conversion of isochorismate to 2-succinyl-5-enolpyruvyl-6-hydroxy-3-cyclohexene-1-carboxylate (SEPHCHC, C), with DHNA acting as a non-competitive inhibitor(19, 20), and the SEPHCHC (C) production rate can be expressed as:

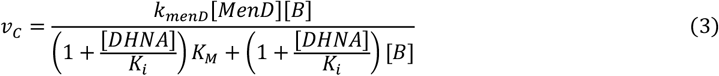

Other enzymes involved, namely MenH, MenD, MenE, MenB, and MenI are modeled using Michaelis-Menten kinetics, and their production rates can be expressed as:

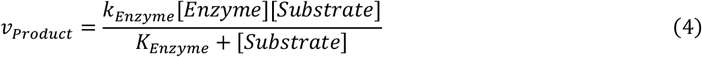

In the first model, we set the chorismate concentration as a constant. The DHNA diffusion through the membrane occurs at rate *k*_*DHNA*_The dynamic equations are:

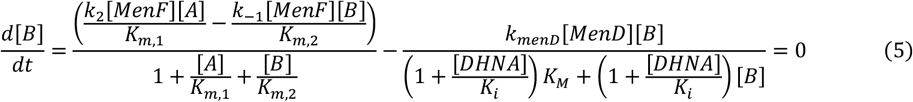

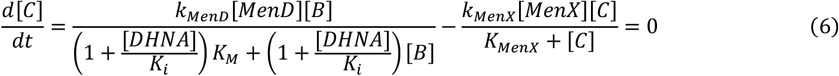

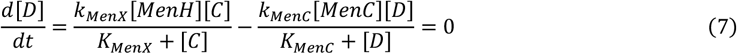

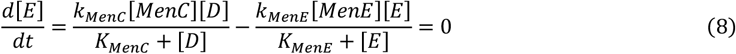

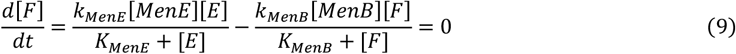

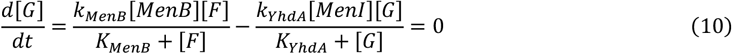

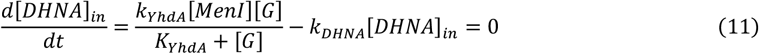

When extracellular DHNA concentration [*DHNA*]_*ex*_ is smaller than the intracellular DHNA concentration, [*DHNA*]_*in*_, the increasing rate of *[DHNA]*_*ex*_ can be calculated as:

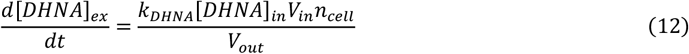

Where *V*_*in*_ is the cell volume, *n*_*cell*_ is the number of cells, *V*_*out*_ is the assay volume. Because the *V*_*in*_, *n*_*cell*_, and *V*_*out*_ are fixed in our experiment, the 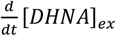 is proportional to [*DHNA*]_*in*_.

By linking these equations, we observe that the reaction flow is primarily regulated by the enzymatic activities of MenF and MenD. Changes in the concentrations of MenH, MenC, MenE, MenB, and MenI are balanced by their respective substrates, leading to a consistent reaction flow across the pathway. Consequently, we simplify the model as follows:

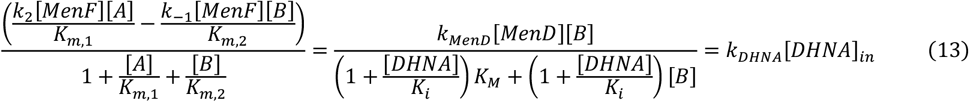

These simplifications highlight the critical roles of MenF and MenD in regulating the pathway’s throughput and underscore the robustness of our model in identifying the rate-limiting steps in DHNA biosynthesis.

##### 1.1 Boundary Conditions Analysis

To depict the scenarios under which the DHNA concentration is regulated predominantly by MenF or MenD, we evaluated specific boundary conditions. These conditions help to define the dynamic behavior of the pathway under various substrate saturations and enzyme concentrations.

###### 1.1.1 Low DHNA Concentration

When DHNA concentration is negligible relative to its inhibition constant 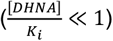, the system simplifies to:

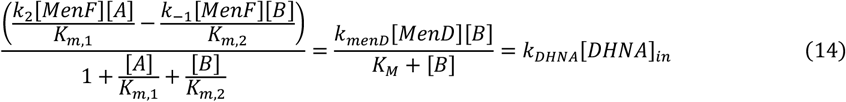

###### 1.1.1.1 High Substrate Conversion Rate ([*B*] ≫ *K*_*M*_)

Under conditions where the intermediate substrate concentration significantly exceeds its Michaelis constant, DHNA concentration becomes regulated solely by MenD, determined by:

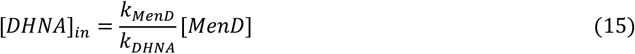

The critical ratio of MenD to MenF that depicts the dominance of MenD in the reaction flux can be described by the following boundary condition:

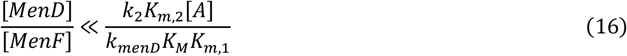

###### 1.1.1.2 When [*B*] ≪ *K*_*m,2*_

DHNA concentration is primarily influenced by MenF:

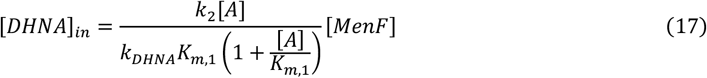

The critical ratio of MenD to MenF that depicts the dominance of MenF in the reaction flux can be described by the following boundary condition:

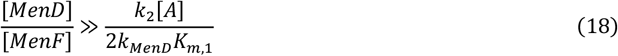

###### 1.1.2 High DHNA Concentration

When DHNA concentration is significantly larger than its inhibition constant 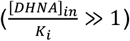, the system simplifies to:

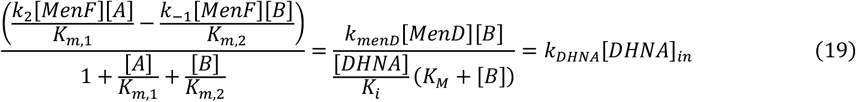

###### 1.1.2.1. When [*B*] ≫ *K*_*M*_, intracellular DHNA concentration becomes regulated solely by MenD

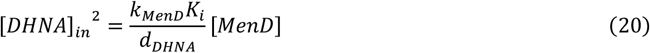

The critical ratio of [*MenD*] to [*MenF*]^2^ that depicts the dominance of MenD in the reaction flux can be described by the following boundary condition:

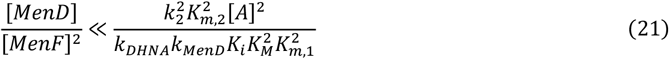

###### 1.1.2.2 When [*B*] ≪ *K*_*m,2*_, DHNA concentration becomes regulated solely by MenF

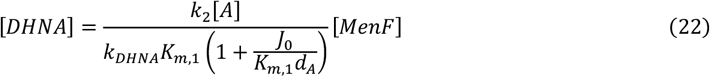

The critical ratio of MenD to MenF^2^ that depicts the dominance of MenF in the reaction flux can be described by the following boundary condition:

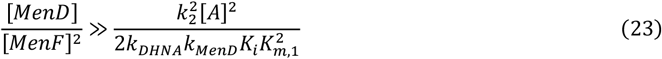

Although this model could explain the conditions where MenD or MenF become saturated and cannot regulate the DHNA synthesis, it cannot explain why perturbation of MenFD does not affect DHNA production. The saturation in MenFD catalysis suggests that the DHNA synthesis is also limited by a bottleneck at the chorismate production.

#### 2. Model 2

Assuming a constant chorismate production rate *J*_0_ and a cumulative chorismite consumption rate *k*_*<*_ by other pathways, the differential equation for chorismite becomes:

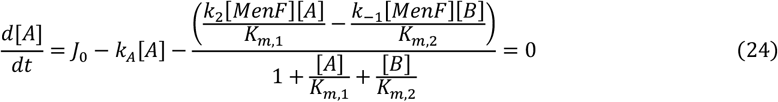

The model becomes:

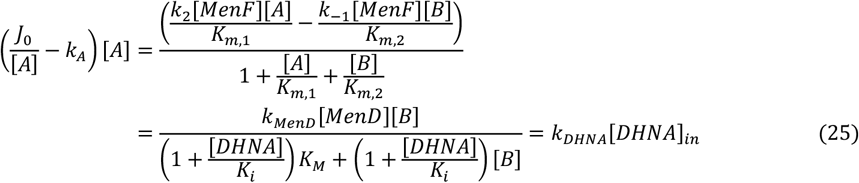

##### 2.1 Substrate-limited Condition

When the pathway for DHNA synthesis is dominant, such that the initial rate of substrate conversion 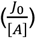 significantly exceeds the reaction rate of the other pathways (*k*_*A*_), the DHNA concentration reaches a maximum, calculated as:

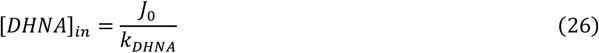

Which is the maximum DHNA concentration we can get.

##### 2.2 Regulation Under Low Substrate Utilization

In contrast, when the pathway utilizes a minimal amount of chorismate compared to other pathways, chorismate concentration is described by:

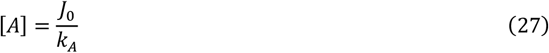

Under this condition, the model can be integrated into model 1 by replacing [*A*] with 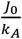 in equation (13).

#### 3. Numerical simulation

To simulate the steady-state concentration of DHNA with given MenF and MenD concentrations, we use the following parameters from the literature:

| Parameter | Value | Organism | Condition | References |
| --- | --- | --- | --- | --- |
| $K_{m,1}$ | 166.9 $\mu M$ | *Escherichia coli* | pH=7.5, 37°C | (43) |
| $K_{m,2}$ | 119 $\mu M$ | | | |
| $k_2$ | 144.9 $min^{-1}$ | | | |
| $k_{-1}$ | 3.8 $min^{-1}$ | | | |
| $K_M$ | 53 nM | *Escherichia coli* | pH=7.8, 22.5°C | (47) |
| $k_{MenD}$ | 3.0 $min^{-1}$ | | | |
| $K_i$ | 53 nM | *Mycobacterium tuberculosis* | pH=8, 20°C | (20) |

The following are parameters used in our assay:

| Parameter | Value |
| --- | --- |
| $V_{in}$ | 1 $\mu m^3$ |
| $n_{cell}$ | $8.55 \times 10^7$ |
| $V_{out}$ | $4.5 \times 10^{11} \mu m^3$ |

For the diffusion rate of DHNA, assuming no external DHNA, the diffusion rate *k*_*DHNA*_ was calculated based on the cell’s surface area (*SA*) and permeability (*α*):

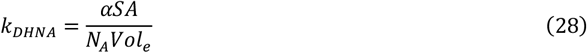

We used the permeability parameter αof small molecules(48). α = 10^−6^ *cm*/*s*. For a spheroidal cell with a radius of 0.5 *μm*, we have:

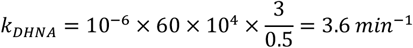

Based on our experimental results in calculating the extracellular DHNA accumulation rate (**Fig. S2**), we hypothesized that the DHNA synthesis had not reached its steady state within 0.5 hours. Thus, the [*DHNA*]_*ex*_ measured after 2 hours of incubation is equivalent to DHNA synthesis at its steady-state rate for 1.5 hours. Using Equations 12, 25, we have:

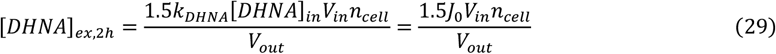

The maximum [*DHNA*]_*ex*_ we observed was 29.51 *nM*. Therefore, we estimated the chorismate synthesis rate *J*_0_ as:

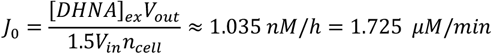

The reaction rate for other pathways was assumed as:

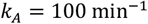

Which is comparable to the k values we used in the DHNA synthesis pathway.

We set the concentration of chorismate in the old model as 1.725 × 10^−2^*μM* for consistency. Using MATLAB, we performed simulations to determine the steady-state concentration of DHNA based on the aforementioned parameters.

## Author Contributions

Conceptualization: S.L., J.Z., C.M.A-F, O.A.I.; Methodology: S.L., J.Z.; Investigation: S.L., J.Z.; Visualization: S.L., J.Z.; Funding acquisition: C.M.A-F, O.A.I.; Supervision: C.M.A-F, O.A.I.; Writing – original draft: S.L., J.Z.; Writing – review & editing: S.L., J.Z., C.M.A-F, O.A.I.

## Acknowledgments

We thank Prof. Maria Marco (UC Davis) for kindly providing the parental strain *L. lactis* subsp. *lactis* KF147. This work was supported by the Cancer Prevention and Research Institute of Texas award # RR190063 and Rice Synthetic Biology Institute.

## Supplementary Figures

**Figure S1:**
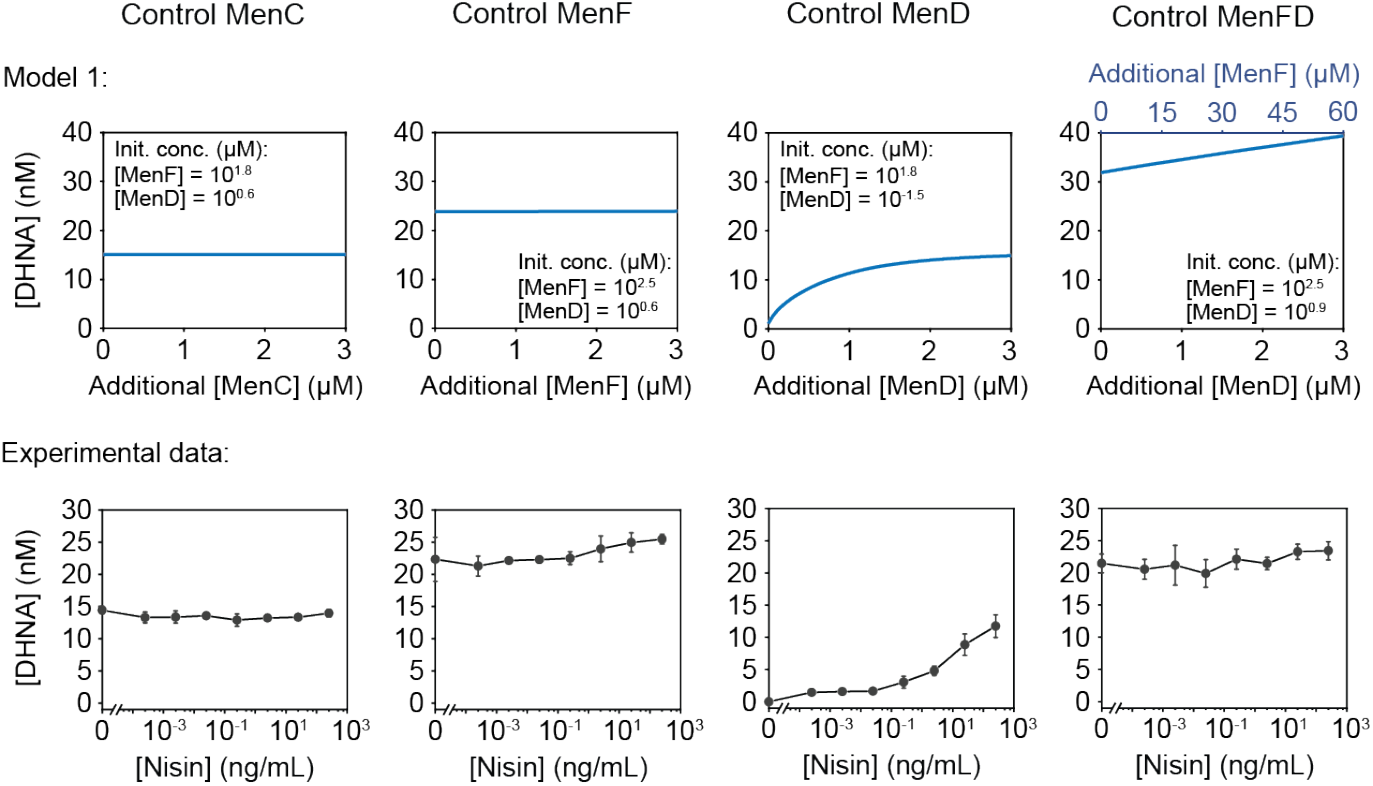
Model 1 correctly predicts DHNA concentrations in response to the perturbations of MenC, MenF, and MenD, but fails when applied to MenFD. The top row represents the predicted changes in DHNA concentrations in response to perturbations of MenC, MenF, and MenD, and MenFD, and the bottom row shows the experimental results. All experimental data represent mean ± 1 s.d. of n=3 biological replicates.

**Figure S2:**
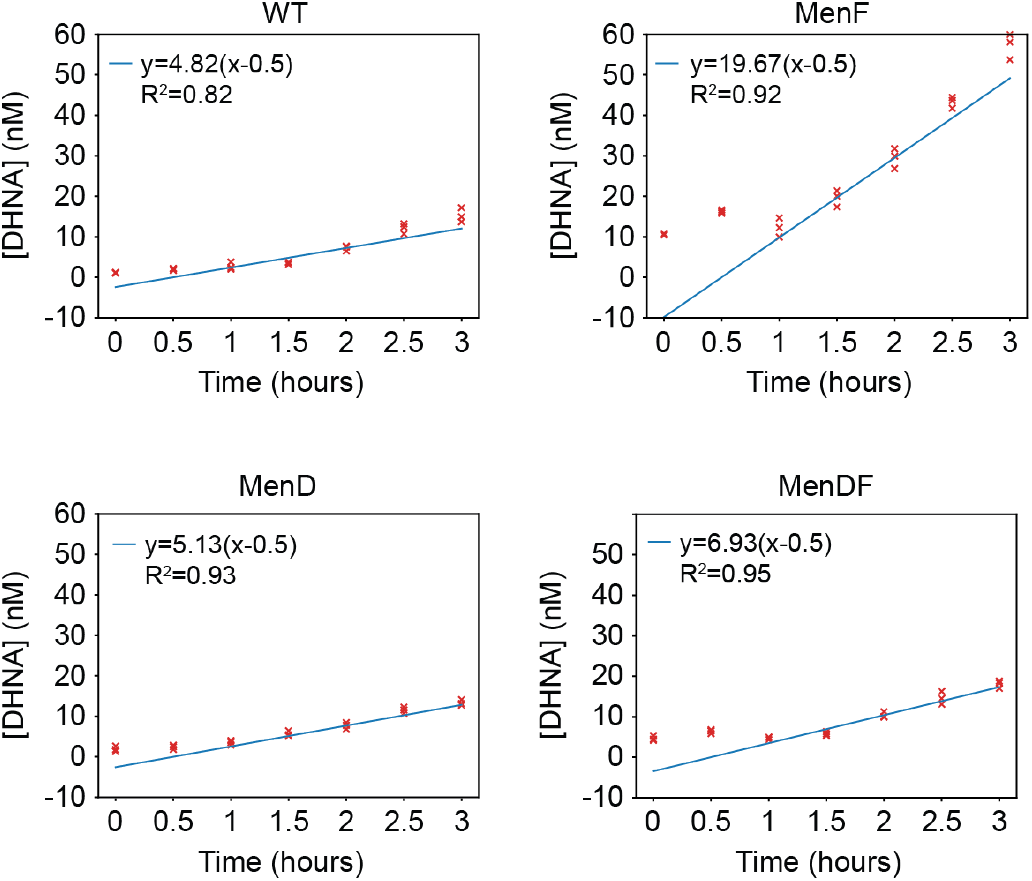
Determination of extracellular DHNA accumulation rate. The supernatants of *L. lactis* were collected at the indicated time points, and the extracellular DHNA concentrations were determined using the EET-based biosensing system. For calculating the DHNA accumulation rate, the linear regression function y=a(x-0.5) was fitted to the experimental data (mean of the three biological replicates) from 1 to 3 hours. The 0 and 0.5 hours data were omitted because we hypothesized that the DHNA outflux had not reached its steady-state rate within 0.5 hours. The MenF, MenD, and MenDF expressions were induced with 25 ng/mL nisin.

## Supplementary Tables

**Table S1.**
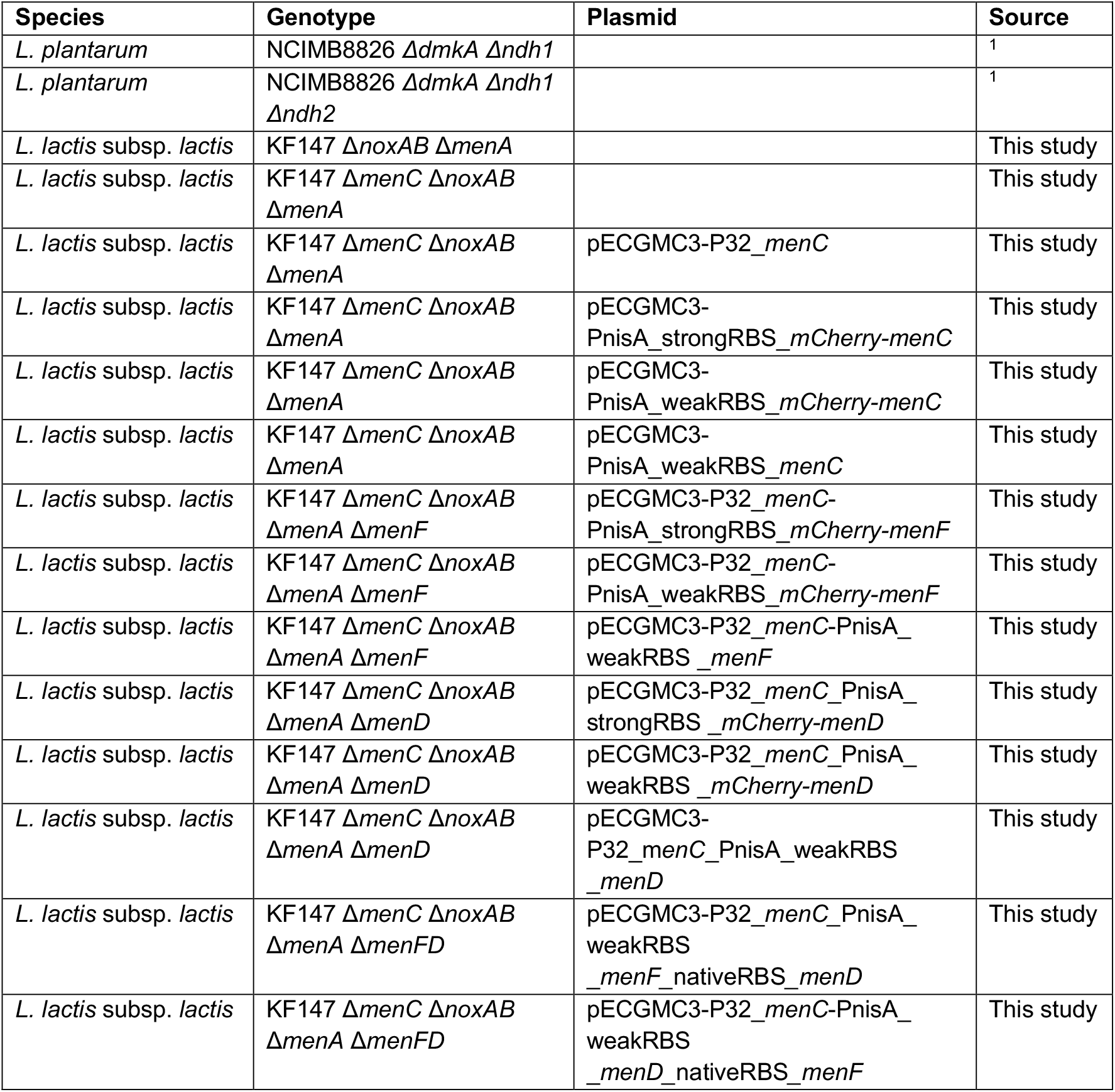
Strains used in this study.

**Table S2.** Recipe of mannitol-MRS (mMRS)

| Component | Final concentration (g/L) |
| --- | --- |
| Protease Peptone #3 | 10 |
| Tween80 | 1 |
| Mannitol | 10 |
| Yeast Extract | 5 |
| Potassium Phosphate dibasic | 2 |
| Sodium acetate trihydrate | 8.3 |
| ammonium citrate tribasic | 2.15 |
| Magnesium sulfate anhydrous | 0.1 |
| Manganese sulfate monohydrate | 0.05 |

**Table S3.** Recipe of mannitol-chemically defined medium (mCDM)

| Component | Final concentration (g/L) |
| --- | --- |
| **Buffers and salts** |  |
| MOPS (3-(N-morpholino)propanesulfonic acid) | 8.371 |
| K<sub>2</sub>HPO<sub>4</sub> | 0.871 |
| NH<sub>4</sub>Cl | 1.070 |
| Na<sub>2</sub>SO<sub>4</sub> | 1.420 |
| **Carbon source** |  |
| Mannitol | 10 |
| **Metals** |  |
| MgCl<sub>2</sub> * 6H<sub>2</sub>O | 0.203 |
| MnCl<sub>2</sub> * 4H<sub>2</sub>O | 0.01 |
| FeSO<sub>4</sub> * 7H<sub>2</sub>O | 0.014 |
| **Amino acids** |  |
| Casamino acids | 1.500 |
| Cysteine-HCl * H<sub>2</sub>O | 0.145 |
| Tryptophan | 0.050 |
| **Wolfe's Vitamins** |  |
| Pyridoxine HCl | 0.001 |
| Thiamine HCl | 0.0005 |
| Riboflavin | 0.0005 |
| Nicotinic acid | 0.0005 |
| Calcium D-(+)-pantothenate | 0.0005 |
| *p*-Aminobenzoic acid | 0.0005 |
| Thioctic acid (α-Lipoic acid) | 0.0005 |
| Biotin | 0.0002 |
| Folic acid | 0.0002 |
| Vitamin B12 | 0.00001 |
| **Wolfe's Minerals** |  |
| Nitrilotriacetic acid (NTA) | 0.15 |
| MgSO<sub>4</sub> * 7H<sub>2</sub>O | 0.3 |
| MnSO<sub>4</sub> * H<sub>2</sub>O | 0.05 |
| NaCl | 0.1 |
| FeSO<sub>4</sub> * 7H<sub>2</sub>O | 0.01 |
| CoCl<sub>2</sub> * 6H<sub>2</sub>O | 0.01 |
| CaCl<sub>2</sub> | 0.01 |
| ZnSO<sub>4</sub> * 7H<sub>2</sub>O | 0.01 |
| CuSO<sub>4</sub> * 5H<sub>2</sub>O | 0.001 |
| AlK(SO)<sub>4</sub> * 12H<sub>2</sub>O | 0.001 |
| H<sub>3</sub>BO<sub>3</sub> | 0.001 |
| Na<sub>2</sub>MoO<sub>4</sub> * 2H<sub>2</sub>O | 0.001 |

## References

1. T. Franza, P. Gaudu, Quinones: more than electron shuttles. Research in Microbiology 173, 103953 (2022).

2. D. K. Newman, R. Kolter, A role for excreted quinones in extracellular electron transfer. Nature 405, 94–97 (2000).

3. S. H. Light, et al., A flavin-based extracellular electron transfer mechanism in diverse Gram-positive bacteria. Nature 562, 140–144 (2018).

4. E. T. Stevens, et al., Lactiplantibacillus plantarum uses ecologically relevant, exogenous quinones for extracellular electron transfer. mBio 14, e02234–23 (2023).

5. E. Mevers, et al., An elusive electron shuttle from a facultative anaerobe. eLife 8, e48054 (2019).

6. A. F. Alvarez, C. Rodriguez, D. Georgellis, Ubiquinone and Menaquinone Electron Carriers Represent the Yin and Yang in the Redox Regulation of the ArcB Sensor Kinase. J Bacteriol 195, 3054–3061 (2013).

7. J. Madeo, A. Zubair, F. Marianne, A review on the role of quinones in renal disorders. Springerplus 2, 139 (2013).

8. J. W. J. Beulens, et al., The role of menaquinones (vitamin K2) in human health. Br J Nutr 110, 1357– 1368 (2013).

9. B. Walther, J. P. Karl, S. L. Booth, P. Boyaval, Menaquinones, Bacteria, and the Food Supply: The Relevance of Dairy and Fermented Food Products to Vitamin K Requirements123. Adv Nutr 4, 463– 473 (2013).

10. E. Manoury, K. Jourdon, P. Boyaval, P. Fourcassié, Quantitative measurement of vitamin K2 (menaquinones) in various fermented dairy products using a reliable high-performance liquid chromatography method. Journal of Dairy Science 96, 1335–1346 (2013).

11. M.-J. Kang, K.-R. Baek, Y.-R. Lee, G.-H. Kim, S.-O. Seo, Production of Vitamin K by Wild-Type and Engineered Microorganisms. Microorganisms 10, 554 (2022).

12. Z. Zhang, L. Liu, C. Liu, Y. Sun, D. Zhang, New aspects of microbial vitamin K2 production by expanding the product spectrum. Microb Cell Fact 20, 1–12 (2021).

13. M. K. Kong, P. C. Lee, Metabolic engineering of menaquinone-8 pathway of Escherichia coli as a microbial platform for vitamin K production. Biotechnol. Bioeng. 108, 1997–2002 (2011).

14. S. Yang, et al., Modular Pathway Engineering of Bacillus subtilis To Promote De Novo Biosynthesis of Menaquinone-7. ACS Synth. Biol. 8, 70–81 (2019).

15. S. Cui, et al., Engineering a Bifunctional Phr60-Rap60-Spo0A Quorum-Sensing Molecular Switch for Dynamic Fine-Tuning of Menaquinone-7 Synthesis in Bacillus subtilis. ACS Synth. Biol. 8, 1826– 1837 (2019).

16. C. A. Bøe, H. Holo, Engineering Lactococcus lactis for Increased Vitamin K2 Production. Front Bioeng Biotechnol 8, 191 (2020).

17. J.-Z. Xu, W.-L. Yan, W.-G. Zhang, Enhancing menaquinone-7 production in recombinant Bacillus amyloliquefaciens by metabolic pathway engineering. RSC Adv. 7, 28527–28534 (2017).

18. Q. Gao, et al., Highly Efficient Production of Menaquinone-7 from Glucose by Metabolically Engineered Escherichia coli. ACS Synth. Biol. 10, 756–765 (2021).

19. T. Stanborough, et al., Allosteric inhibition of Staphylococcus aureus MenD by 1,4-dihydroxy naphthoic acid: a feedback inhibition mechanism of the menaquinone biosynthesis pathway. Philosophical Transactions of the Royal Society B: Biological Sciences 378, 20220035 (2023).

20. G. Bashiri, et al., Allosteric regulation of menaquinone (vitamin K2) biosynthesis in the human pathogen Mycobacterium tuberculosis. J Biol Chem 295, 3759–3770 (2020).

21. Y. Tani, S. Asahi, H. Yamada, Menaquinone (vitamin K2)-6 production by mutants of Flavobacterium meningosepticum. J Nutr Sci Vitaminol (Tokyo) 32, 137–145 (1986).

22. S. Tachon, J. B. Brandsma, M. Yvon, NoxE NADH Oxidase and the Electron Transport Chain Are Responsible for the Ability of Lactococcus lactis To Decrease the Redox Potential of Milk. Appl Environ Microbiol 76, 1311–1319 (2010).

23. K. Isawa, et al., Isolation and Identification of a New Bifidogenic Growth Stimulator Produced by Propionibacterium freudenreichii ET-3. Bioscience, Biotechnology, and Biochemistry 66, 679–681 (2002).

24. K. Furuichi, Y. Katakura, K. Ninomiya, S. Shioya, Enhancement of 1,4-Dihydroxy-2-Naphthoic Acid Production by Propionibacterium freudenreichii ET-3 Fed-Batch Culture. Appl Environ Microbiol 73, 3137–3143 (2007).

25. J.-E. Eom, S.-C. Kwon, G.-S. Moon, Detection of 1,4-Dihydroxy-2-Naphthoic Acid from Commercial Makgeolli Products. Prev Nutr Food Sci 17, 83–86 (2012).

26. J. Takebayashi, J. Nagata, K. Yamada, Improved Analytical Precision of 1,4-Dihydroxy-2-naphthoic Acid by High Performance Liquid Chromatography Using Dithiothreitol as Mobile Phase Additive. FSTR 14, 509–512 (2008).

27. S. Tejedor-Sanz, et al., Extracellular electron transfer increases fermentation in lactic acid bacteria via a hybrid metabolism. eLife 11, e70684 (2022).

28. J. G. Tolar, S. Li, C. M. Ajo-Franklin, The Differing Roles of Flavins and Quinones in Extracellular Electron Transfer in Lactiplantibacillus plantarum. Applied and Environmental Microbiology 89, e01313–22 (2022).

29. S. Li, C. De Groote Tavares, J. G. Tolar, C. M. Ajo-Franklin, Selective bioelectronic sensing of pharmacologically relevant quinones using extracellular electron transfer in Lactiplantibacillus plantarum. Biosensors and Bioelectronics 243, 115762 (2024).

30. R. Meganathan, Biosynthesis of menaquinone (vitamin K2) and ubiquinone (coenzyme Q): a perspective on enzymatic mechanisms. Vitam Horm 61, 173–218 (2001).

31. P. G. de Ruyter, O. P. Kuipers, W. M. de Vos, Controlled gene expression systems for Lactococcus lactis with the food-grade inducer nisin. Appl Environ Microbiol 62, 3662–3667 (1996).

32. J. M. Johnston, E. M. Bulloch, Advances in menaquinone biosynthesis: sublocalisation and allosteric regulation. Current Opinion in Structural Biology 65, 33–41 (2020).

33. F. Hubrich, M. Müller, J. N. Andexer, Chorismate- and isochorismate converting enzymes: versatile catalysts acting on an important metabolic node. Chemical Communications 57, 2441–2463 (2021).

34. H. N. Lim, Y. Lee, R. Hussein, Fundamental relationship between operon organization and gene expression. Proceedings of the National Academy of Sciences 108, 10626–10631 (2011).

35. M. N. Price, A. P. Arkin, E. J. Alm, The Life-Cycle of Operons. PLOS Genetics 2, e96 (2006).

36. L. Gu, et al., Rewiring the respiratory pathway of Lactococcus lactis to enhance extracellular electron transfer. Microbial Biotechnology 16, 1277–1292 (2023).

37. L. Rezaïki, G. Lamberet, A. Derré, A. Gruss, P. Gaudu, Lactococcus lactis produces short-chain quinones that cross-feed Group B Streptococcus to activate respiration growth. Mol. Microbiol. 67, 947–957 (2008).

38. S. Tachon, J. B. Brandsma, M. Yvon, NoxE NADH Oxidase and the Electron Transport Chain Are Responsible for the Ability of Lactococcus lactis To Decrease the Redox Potential of Milk. Appl Environ Microbiol 76, 1311–1319 (2010).

39. C. A. Abbas, A. A. Sibirny, Genetic Control of Biosynthesis and Transport of Riboflavin and Flavin Nucleotides and Construction of Robust Biotechnological Producers. Microbiol Mol Biol Rev 75, 321–360 (2011).

40. L. S. Pierson, E. A. Pierson, Metabolism and function of phenazines in bacteria: impacts on the behavior of bacteria in the environment and biotechnological processes. Appl Microbiol Biotechnol 86, 1659–1670 (2010).

41. D. Cavanagh, G. F. Fitzgerald, O. McAuliffe, From field to fermentation: The origins of Lactococcus lactis and its domestication to the dairy environment. Food Microbiology 47, 45–61 (2015).

42. X. Qin, H. W. Taber, Transcriptional regulation of the Bacillus subtilis menp1 promoter. Journal of Bacteriology 178, 705–713 (1996).

43. C. Dahm, R. Müller, G. Schulte, K. Schmidt, E. Leistner, The role of isochorismate hydroxymutase genes entC and menF in enterobactin and menaquinone biosynthesis in Escherichia coli. Biochimica et Biophysica Acta (BBA) - General Subjects 1425, 377–386 (1998).

44. Y. Tsukamoto, M. Kasai, H. Kakuda, Construction of a Bacillus subtilis (natto) with high productivity of vitamin K2 (menaquinone-7) by analog resistance. Biosci Biotechnol Biochem 65, 2007–2015 (2001).

45. L. Leloup, S. D. Ehrlich, M. Zagorec, F. Morel-Deville, Single-crossover integration in the Lactobacillus sake chromosome and insertional inactivation of the ptsI and lacL genes. Appl Environ Microbiol 63, 2117–2123 (1997).

46. C. Engler, R. Kandzia, S. Marillonnet, A One Pot, One Step, Precision Cloning Method with High Throughput Capability. PLoS ONE 3, e3647 (2008).

47. M. Jiang, et al., Menaquinone Biosynthesis in Escherichia coli : Identification of 2-Succinyl-5-enolpyruvyl-6-hydroxy-3-cyclohexene-1-carboxylate as a Novel Intermediate and Re-Evaluation of MenD Activity. Biochemistry 46, 10979–10989 (2007).

48. N. J. Yang, M. J. Hinner, Getting Across the Cell Membrane: An Overview for Small Molecules, Peptides, and Proteins. Methods Mol Biol 1266, 29–53 (2015).

## Supplementary reference

1. Li, S., De Groote Tavares, C., Tolar, J.G., and Ajo-Franklin, C.M. (2024). Selective bioelectronic sensing of pharmacologically relevant quinones using extracellular electron transfer in Lactiplantibacillus plantarum. Biosensors and Bioelectronics 243, 115762. 10.1016/j.bios.2023.115762.

